# Pre-zygotic mate selection affects progeny fitness and is only partially correlated with the expression of *Na*S-like RNases

**DOI:** 10.1101/2023.09.27.559675

**Authors:** Patrycja Baraniecka, Wibke Seibt, Karin Groten, Danny Kessler, Erica McGale, Klaus Gase, Ian T. Baldwin, John R. Pannell

## Abstract

- *Nicotiana attenuata* styles preferentially select pollen from accessions with corresponding expression patterns of *Na*S-like-RNases (SLRs), and the post-pollination ethylene burst (PPEB) is an accurate predictor of seed siring success. However, the ecological consequences of mate selection, its effect on the progeny, and the role of SLRs in the control of ethylene signaling are still not well understood.
- We explored the link between the magnitude of the ethylene burst and transcript and protein abundance of the SLRs in a set of recombinant inbred lines (RILs) and investigated the fitness consequences of mate selection for the next generation. Genome Wide Association Study (GWAS) identified novel candidate genes potentially involved in the control of mate selection.
- We found that high levels of PPEB are associated with the absence of SLR2 but not with the expression of SLR1 in most of the tested RILs. Maternal genotypes that favor certain pollen produce offspring with longer roots when pollinated with these donors, but the selection for beneficial traits is abolished when the maternal genotype selects only against certain pollen donors.
- We conclude that mate selection mechanisms affect the offspring number and performance in ways that might be adaptive.

## Introduction

Flowering plants have evolved a variety of different mating systems that differ in mechanism and pollen preference (Pannell & Voillemot, 2017; Whitehead *et al*., 2018). One of the best characterized examples is the rejection of self pollen in self-incompatible (SI) species, ensuring the genetic diversity of the progeny (Muñoz-Sanz *et al*., 2020; Goring *et al*., 2022). All three SI systems known to date utilize the highly polymorphic female- and male-specific S-determinants, but act via different rejection mechanisms. In Brassicaceae and *Papaver,* the rejection of the incompatible pollen tubes of self pollen is based on self-recognition, while the S-RNase based system in Solanaceae employs a non-self recognition mechanism (Fujii *et al*., 2016). In self-incompatible tobacco and other Solanaceous species, the female S-determinant encodes pistil-specific S-RNases that inhibit the growth of incompatible pollen tubes of self pollen by degrading their RNA via cytotoxic activity (McClure *et al*., 1990). The pollen-specific S-determinant encodes multiple S-locus F-box (SLF) genes (Sun *et al*., 2018) that act collaboratively to neutralize multiple allelic variants of S-RNases (Kubo *et al*., 2010). The SI-specific factors such as S-RNases, SLF proteins, HT-B and Cullin have also been shown to be involved in interspecific interaction of unilateral incompatibility (UI; Li & Chetelat, 2015). Interspecific pollen rejection involves the rejection of pollen from self-compatible species by self-incompatible females, but not *vice versa* (Hancock *et al*., 2003).

In addition to the well described SI and UI systems that both utilize the S-RNases, polyandrous pre-zygotic mate selection was recently identified in *Nicotiana attenuata* (Bhattacharya & Baldwin, 2012; Guo *et al*., 2019). In this self-compatible (SC) diploid wild tobacco species, every cross is accepted and self-fertilization is common. However, in genetically diverse natural populations with high outcrossing rates (Sime & Baldwin, 2003), certain pollen donors are consistently favored over others in mixed pollinations. In particular, self pollen is strongly favored in mixed, binary pollinations with equal numbers of self and non-self pollen grains, despite no differences in seed siring ability for both pollen donors (Bhattacharya & Baldwin, 2012). The study of the molecular basis of pre-zygotic mate selection revealed the involvement of two *NaS-like- RNases* (SLRs) and six *NaSLF-like* genes that are homologues to the self-incompatibility factors in the Solanaceae. In their mixed pollination experiment with 14 different non-self pollen donors, Guo and colleagues (2019) showed that accessions of *N. attenuata* that express SLRs favor mates with a corresponding SLR expression pattern (Fig. 1a,b). Briefly, lines expressing both SLRs were found to strongly favor pollen donors with the same SLR expression pattern (Fig. 1a,b, in grey), while natural accessions and transgenic lines expressing only SLR1 strongly disfavored pollen donors that do not express SLRs (Fig. 1a,b, in pink). No significant differences in the seed siring success among the 14 pollen donors were observed for natural and transgenic lines that completely lack the stylar expression of SLRs (Fig. 1a,b, in blue; Guo *et al*., 2019). Mate selection in *N. attenuata* occurs in the upper part of the style, and pollination with favored pollen results in a predictably higher post-pollination ethylene burst (PPEB). Importantly, the ability of mate choice is completely lost when the ethylene signaling is disrupted (Bhattacharya & Baldwin, 2012). Previous studies demonstrated that the variation in ethylene burst is associated with the efficiency of pollen tubes to penetrate the pistil and emphasized the pivotal role of ethylene signaling in pollen recognition (De Martinis *et al*., 2002). It has been shown that PPEB promotes pollen tube growth through the stylar transmitting tract (Holden *et al*., 2003; Jia *et al*., 2018) and that it can modify the cell wall structure of growing pollen tubes through the regulation of genes involved in pectin and hemicellulose modifications (Althiab-Almasaud *et al*., 2021).

**Fig. 1:**
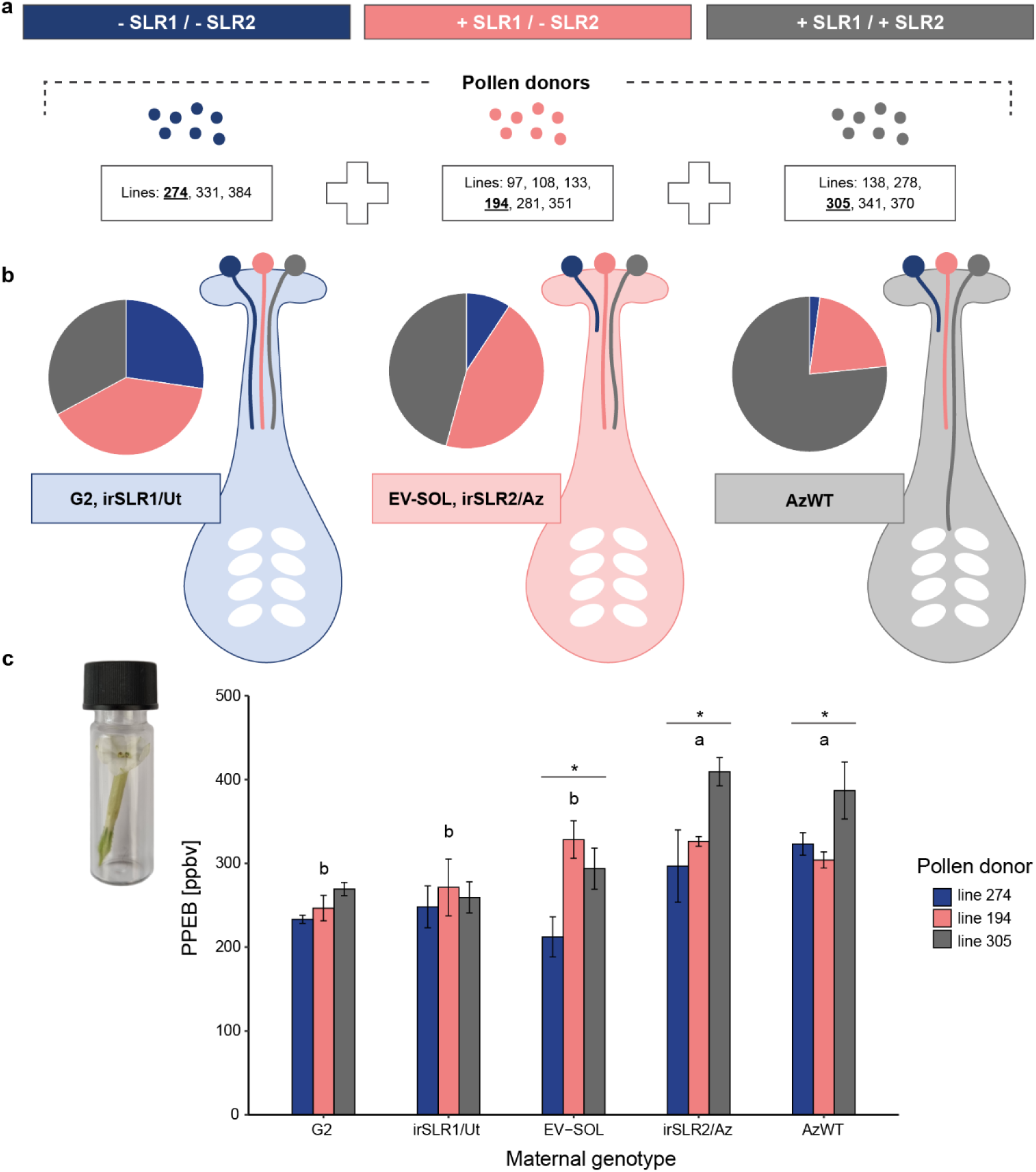
The mate selection patterns in *Nicotiana attenuata* mixed pollination correlate with the expression pattern of the self-incompatibility factors S-like-RNases (SLRs) in the maternal genotypes, but the post-pollination ethylene burst (PPEB) only partially corresponds with this pattern. (a) Pollen donors and (b) maternal genotypes can be divided into three groups depending on the expression of SLRs: lines that do not express any of the two SLRs are shown in blue, lines that express only SLR1 are shown in pink and lines that express both SLRs are shown in grey. The representative lines selected from each SLR expression group for further experiments are underlined. (b) Styles preferentially select pollen from accessions with the corresponding SLR expression pattern. The schematic pie charts represent the percentage of seed siring success by different pollen donors (shown in a) after mixed pollination according to Guo et al. (2019). The colors of slices, schematic styles and pollen grains correspond to different SLR expression groups. Different lengths of pollen tubes in the style correspond to differences in mate selection patterns. (c) PPEB in genotypes shown in (b) after single pollination with a representative pollen donor from each SLR expression group. Data are shown as means ± se, n = 3. The statistical differences were calculated using two-way ANOVA and TukeyHSD post hoc test, different letters indicate significant differences dependent on the maternal genotype, asterisks indicate significant differences dependent on the pollen donor, *P* < 0.05.

The ecological function of pre-zygotic mate selection and the exact role of the SI elements in a fully self-compatible species are still not well understood. One possibility is that the pre-zygotic barriers in the genotypes that express SLRs favor particular mates with traits that confer potential fitness advantages through the production of higher numbers of superior offspring (Bernasconi *et al*., 2004). It has been suggested that the SI elements might have been repurposed during the transition from SI to SC to facilitate mate selection via a mechanism that differs from cytotoxic activity usually associated with the classic SI model (Takayama & Isogai, 2005). Alternatively, the transition to SC could have rendered the SI system non-functional, resulting in gradual pseudogenization of its elements. As a consequence, the accessions expressing the SLRs might be in a transitional stage, with their fitness reduced compared to the accessions that have already lost the SLR expression (Guo *et al*., 2019). In the present study, we show that the mate selection mechanisms have a quantifiable effect on the fitness of the progeny and might therefore sometimes, be adaptive. We also show that high level of PPEB is negatively correlated with the expression of SLR2, but not SLR1 for most of the tested lines. As our results indicate that additional regulatory elements are involved in the mate selection processes, we performed a Genome Wide Association Study (GWAS) on PPEB data from a 26-parent Multiparent Advanced Generation Inter-Cross (MAGIC) population and provide a list of seven novel candidates for further investigation.

## Materials and Methods

### Plant material

*Nicotiana attenuata* Torrey Ex Watson Utah (UtWT) and Arizona (AzWT) wild type seeds were originally collected from a large natural population growing near Santa Clara, Utah, USA (Halitschke *et al*., 2000) and a 20-plant population near Flagstaff, Arizona, USA (Glawe *et al*., 2003), respectively. The seeds were inbred for 30 generations for UtWT and for 21 generations for AzWT in the glasshouse. Seeds of the G2 accession were also collected in Utah, USA (Schuman *et al*., 2009). The seeds of natural accessions used as pollen donors: lines 274, 194 and 305 were collected from different populations throughout southwestern US as described previously (Li *et al*., 2015; Guo *et al*., 2019) and inbred for one generation in the glasshouse. Selected recombinant inbred lines (RILs) of a 26-parent MAGIC population (Ray *et al*., 2019) as well as one replicate of the entire population (650 lines) were used. Additionally, a UtWT inbred line silenced in the expression of *Na*S-like-RNase1 (irSLR1/UT, pSOL8SRN1, A-17-091-7), and an AzWT inbred line silenced in the expression of *Na*S-like-RNase2 (irSLR2/AZ, pSOL8SRN2, A- 17-059-6; Guo *et al*., 2019) were used. An empty vector (EV-SOL, pSOL3NC, A-04-266-3) was used as an additional control for the transgenic lines and in the analysis of the offspring, as the mate selection pattern for this specific line has been characterized previously (Bhattacharya & Baldwin, 2012; Guo *et al*., 2019). EV-SOL was produced from UtWT inbred line and hence it is isogenic except for a single T-DNA insertion and potential uncharacterized changes that might have occurred during the transformation process. It has been fully characterized previously (Bubner *et al*., 2006) and is routinely used in the department.

### Growth conditions

For the single pollination experiment and the ethylene measurements as well as for the selected MAGIC RIL lines, seeds were germinated as described previously (Krügel *et al*., 2002) and grown in the glasshouse under long day conditions (26 ± 1°C; 16 h : 8 h, light : dark). For the seedling phenotyping experiment, seeds were germinated on square (12 cm x 12 cm) plates with GB5 medium. The plates were placed vertically in a Percival growth chamber (CLF PlantClimatics GmbH, Wertingen, Germany) for 14 days (26 ± 1°C; 16 h : 8h, light : dark, 75% light intensity). The seeds of the MAGIC RIL lines grown for the GWAS experiment were first treated with 50x diluted liquid smoke solution (House of Herbs, Passaic NJ) and 0.1 M GA3 for 1 h as described by Krügel et al. (2002). Subsequently, the solution was replaced with 100x diluted smoke and the seeds were kept in the fridge overnight before they were directly sown into 4-liter pots and covered with a transparent, plastic cup for 12-16 days until the cotyledons turned to a mat green color. The cup was then removed gradually to allow further growth of the seedlings.

### Single pollination experiment

The natural accessions and transgenic lines used as parental genotypes were selected based on their expression of *Na*S-like-RNases: line 274, which does not express SLR1 or SLR2, line 194, which expresses only SLR1, and line 305, which expresses both SLR1 and SLR2 served as the pollen donors. AzWT served as maternal genotype known to express both SLRs, EV-SOL and irSLR2/Az (AzWT transgenic line silenced in the expression of SLR2) served as maternal genotypes expressing only SLR1, G2 and irSLR1/Ut (UtWT transgenic line silenced in the expression of SLR1) served as maternal genotypes that lack the stylar expression of SLRs (Fig. 1a,b; Guo *et al*., 2019). In the morning before the experiment (06:00 – 08:00 h), mature, open flowers and seed capsules were removed from all the plants. The remaining, immature flowers of the maternal genotypes were antherectomized, as described previously (Kessler *et al*., 2008), to avoid self-pollination. Single pollination was performed in the evening (18:00 – 20:00 h) once all the flowers had opened and the anthers of the pollen donors had matured. The hand pollination was performed by gently rubbing one, freshly matured anther over the stigma. The pollinated flowers were labelled with different colors of threads for each pollen donor gently tied around the pedicel. Depending on the flower availability, four to six flowers uniformly distributed around the plant were pollinated by a single pollen donor. The matured capsules were collected about two weeks later, right before their opening and further dried in silica gel.

### Ethylene measurements

To ensure that the age of flowers used for the measurements was the same across all plants, all mature, open flowers and seed capsules were cut from all parental genotypes either around noon (11:00– 12:00 h) for the next-day morning measurement of the MAGIC RIL lines, or early in the morning (06:00 - 08:00 h) for measurement of the maternal genotypes the same evening (Fig. 1c). Flowers of each RIL line (Fig. 2) were hand-pollinated with UtWT pollen (n = 6), and flowers of the five maternal genotypes (Fig. 1c) were pollinated by lines 274, 194 and 305 (n = 3; see the ‘*Single pollination experiment*’ section). Right after pollination, the flowers were removed from the plant and transferred to a 4 ml glass vial. After 5 h, ethylene was measured from the headspace using a highly sensitive (0.3 ppbv detection limit) ETD-300 Ethylene Detector attached to a VC- 6 Valve Control Box (Sensor Sense, The Netherlands) in the sample mode.

**Fig. 2:**
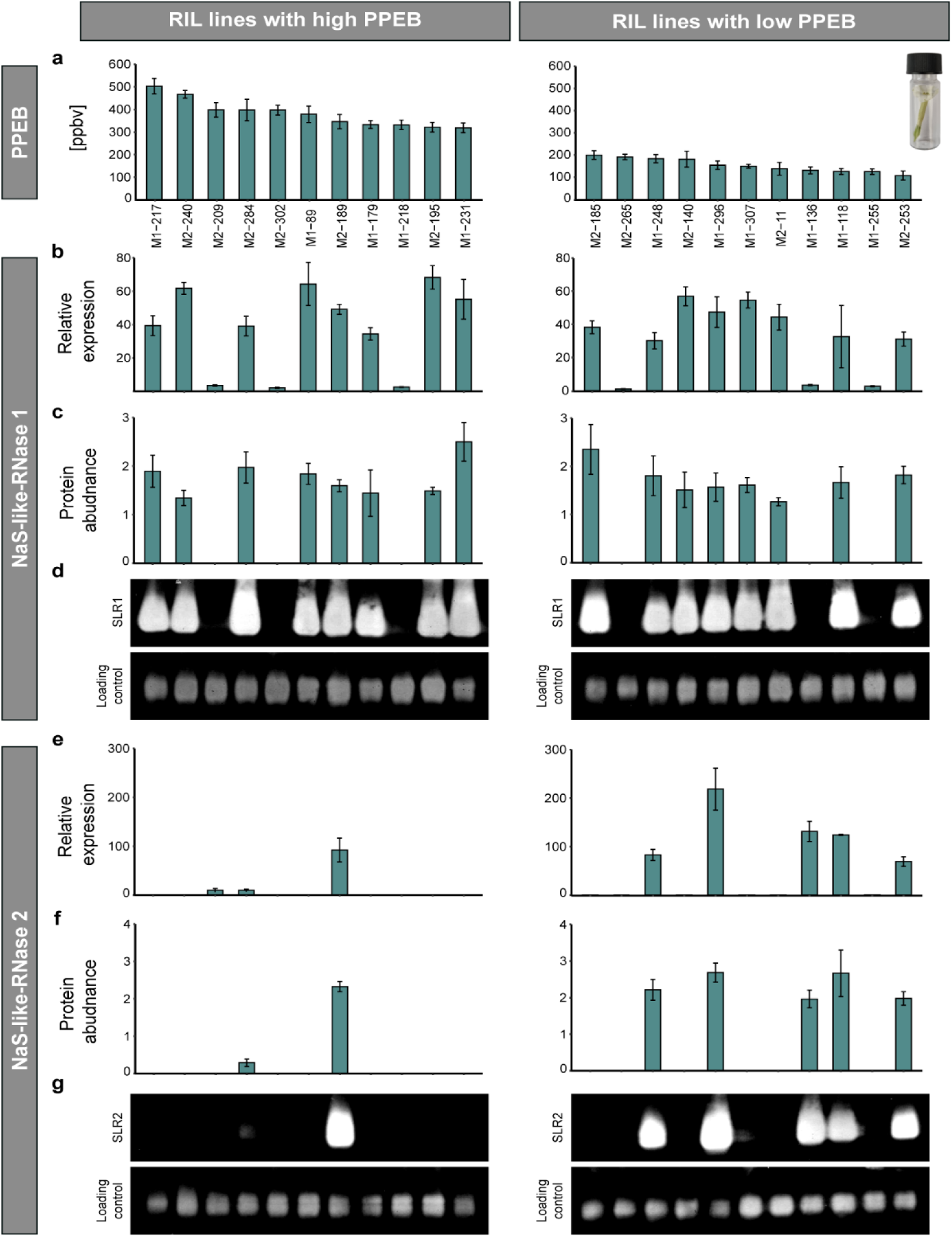
A high magnitude of the post-pollination ethylene burst (PPEB) correlates with the lack of expression of SLR2 in *Nicotiana attenuata*. (a) Quantification of the post-pollination ethylene burst (PPEB) in recombinant inbred lines (RILs) from the 26-parent Multiparent Advanced Generation Inter-Cross (MAGIC) population with extreme phenotypes (high and low) of PPEBs when pollinated with a control, Utah wild type accession (UtWT). Data are shown as means ± se (n = 6). Gene expression of (b) SLR1 and (e) SLR2 measured in unpollinated styles relative to *N. attenuata* elongation factor 1a *Na*EF1a. Protein abundance of (c) SLR1 and (f) SLR2 quantified from the X-ray films shown in (d, g, respectively) based on mean signal intensity relative to an anti-actin loading control using Fiji software. Data are shown as means ± se (n = 5). The brightness and contrast of the western blot results (d, g) were adjusted for imaging purposes, but not before the intensity measurement.

### Quantitative RT-PCR

RNA was extracted from at least five unpollinated styles per replicate (n = 5) using the ‘DNA, RNA, and protein purification’ kit (Macherey-Nagel GmbH & Co. KG, Düren, Germany) according to the manufacturer’s instructions. The RNA samples were further processed using the ‘RNA Clean and Concentrator ™ -5’ kit (Zymo Research, Irvine, CA, USA), following the manufacturer’s protocol. Total RNA was quantified using NanoDrop (Thermo Scientific, Wilmington, DE, USA) and cDNA was synthesized from 500 ng of total RNA using RevertAid H Minus reverse transcriptase and oligo (dT) primer (Fermentas, Vilnius, Lithuania). Quantitative reverse transcription PCR (RT-qPCR) was performed using a Mx3005P PCR cycler (Stratagene, San Diego, CA, USA) and a SYBR green reaction mix with added ROX dye (Eurogentec, Liège, Belgium). The primer sequences used in this study are shown in Table S1. *N. attenuata* elongation factor 1a (*Na*EF1a) was used as an internal control. qRT-PCR data were normalized using the delta-delta-Ct method.

### Western blot analysis

Total protein was extracted from at least five unpollinated styles per replicate (n = 5) using the RB^+^ buffer (1M Tris-HCL, pH 7.8; 5M NaCl; 50% Glycerol; 20% Tween20, β-Me). Briefly, 80 µl of RB^+^ buffer was added to the ground tissue, and the samples were centrifuged twice for 20 min in 4°C at 13.2 krpm. The amount of total protein was quantified by a Bradford assay using Albumin Bovine (BSA, Sigma-Aldrich, Steinheim, Germany) and the Quick Start Bradford 1x Dye Reagent (Bio-Rad Laboratories, Hercules, CA, USA). The light absorbance was measured after 10 min at 595 nm using the Infinite M2000 microplate absorbance reader (TECAN, Grödig, Austria). The samples were then denaturized at 95 °C for 5 min. 15 µg of total protein in 20 µL of sample was loaded on a 12% polyacrylamide gel and run for 40 min at 60 V followed by 90-120 min at 120 V. The protein was transferred onto a PVDF membrane using the Mini Trans-Blot Electrophoretic Transfer Cell (Bio-Rad Laboratories, Hercules, CA, USA) at 320 mA for 90 min. The membrane was blocked overnight in 5% solution of milk powder (Carl Roth GmbH + Co. KG, Karlsruhe, Germany). Subsequently, the membrane was incubated with 1:5000 dilution of the SLR1 specific polyclonal antibody or 1:3000 dilution of the SLR2 specific polyclonal antibody for 1 h. Both antibodies were obtained in rabbits from the synthetically synthesized immunogens based on the protein sequence of SLR1 and SLR2 (GenScript Biotech, Leiden Netherlands). The membranes were then washed four times for 40 min with the 1 x TBST buffer followed by 1 h incubation with commercially available anti-rabbit secondary antibody (Cytvia, Marlborough, MA, USA) and a 1 h wash with 1 x TBST. The protein was detected via chemiluminescence using the ECL™ Western Blotting Detection System (Cytvia, Amersham, UK), and the signal was visualized on CL-XPosure ™ Film (Thermo Scientific, Waltham, MA, USA). The films were developed and fixed using the Readymatic system (Carestream Health, Inc., Belgium). The membranes were then stripped using the 1x Restore Western Blotting Stripping Buffer (Thermo Scientific, Rockford, IL, USA), according to the manufacturer’s instructions, and blocked again in 5% milk solution overnight. Subsequently, they were incubated for 2.5 h with commercially available mouse monoclonal anti-actin antibody specific for plants (Sigma-Aldrich, Saint Louis, USA) diluted 1:500 as a loading control. The membranes were then washed, as described, and incubated with anti-mouse IgG, peroxidase-linked, species-specific secondary antibody for 1 h (Thermo Fisher Scientific) followed by 1 h wash with 1x TBST. The protein was detected via chemiluminescence and the signal was visualized as described.

### Seed count and size measurement

The seeds obtained after the single pollination experiment were first removed from the dried capsule, cleaned from any remaining plant tissue, and weighed. The seeds were then scattered on a square plate and the plate, was scanned using a regular office scanner re-built to allow plate scanning. The pictures were analyzed using Fiji software (version 2.3.0/1.53q) and the particle analysis tool. Seed number and average seed size data were collected and are presented as an average per capsule.

### Seedling phenotyping

Ten seeds from three capsules per cross obtained from the single pollination experiment were germinated on square plates (2 cm from the top of the plate and in 1 cm distance from one another) and grown vertically for 14 days, as described (see the ‘*Growth conditions*’ section). All plates were scanned daily starting three days after sowing (DAS) using the regular office scanner re-built to allow plate scanning to determine the germination date. Once all seeds germinated, the plates were scanned every second day from the top and bottom. The root and shoot biomass of each seedling were collected destructively at the end of the experiment, 14 DAS. Data for root length, root area, hypocotyl length and rosette size were obtained from the pictures using the Fiji software (version 2.3.0/1.53q) for six time points. Representative picture of seedlings from each cross was taken using a regular mobile phone camera (Fig. S3).

### Genome Wide Association Study (GWAS)

Reciprocal hand pollination was performed 650 recombinant inbred lines (RILs) from a 26-parent Multiparent Advanced Generation Inter-Cross (MAGIC) population (Ray *et al*., 2019) using pollen from the UtWT accession, and UtWT was pollinated with pollen from each of the 650 RIL lines from the MAGIC population (n = 1), and the PPEB was measured, as described (see the ‘*Ethylene measurement*’ section). These crosses resulted in two different datasets (Fig. S1). The initial data was examined for potential variation due to the time of pollination or differences in environmental conditions and inconsistent values were removed. GWAS was conducted on 629 datapoints in each dataset using the Genomic Association and Prediction Integrated Tool (GAPIT version 3) package in the R interface (Wang & Zhang, 2021). Generalized Linear Models (GLM) were used for the association analysis. Significance of associations was inferred on the basis of an LOD threshold of 1e-05.

### Statistical analysis

All data were analyzed using R statistical Software version 4.2.1 (R Core Team, 2020) in RStudio version 2022.07.1 + 554 (RStudio Team, 2020). The datasets regarding seed attributes (Fig. 3) and early seedling establishment (Fig. 5) were first tested for the effect of the individual using ANCOVA, intraclass correlation and bootstrapping. Whenever the effect of the individual was significant, the data were fit to a linear mixed-effect model (using the *nlme* package) to account for the nested structure of the data per individual and checked for homoscedasticity and normality through a graphical analysis of residuals (Zuur *et al*., 2009). Otherwise, two-way ANOVAs were applied (using the *emmeans* package; Lenth *et al*., 2019). Multiple comparisons were extracted with a Sidak (for the mixed-effect models) or TukeyHSD (for the ANOVAs) *post-hoc* test. The correlations shown in Fig. 3 and Table S2 were extracted using the Spearman’s rank correlation coefficient.

**Fig. 3:**
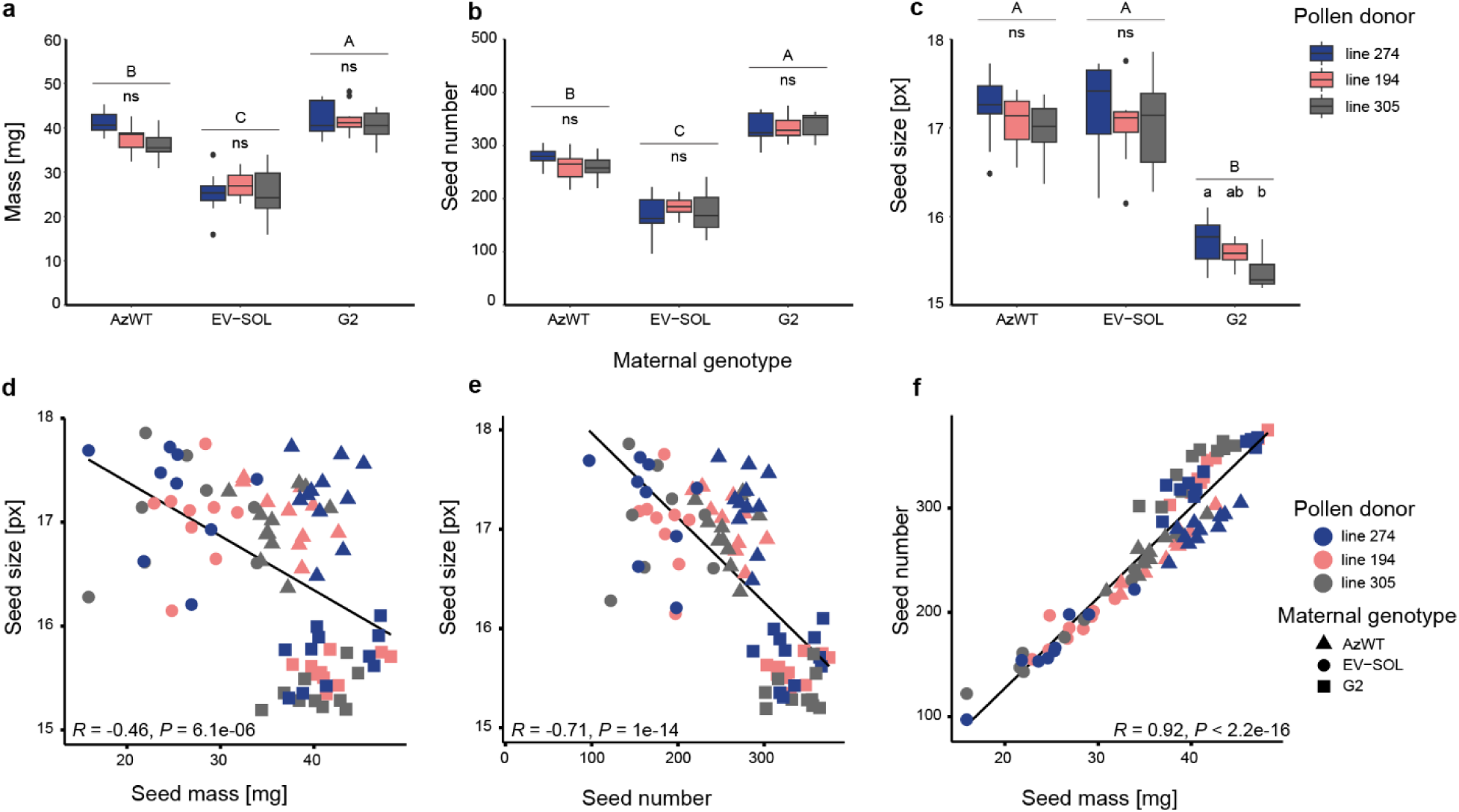
In *Nicotiana attenuata* the tradeoff between seed size, seed mass and number depends on maternal genotype. (a) The seed mass, (b) seed number and (c) average seed size were calculated per capsule produced by different maternal genotypes after single hand pollination with a favored, non-favored or neutral pollen donor. Data are shown as means of up to ten capsules collected from two different plants per mother genotype obtained after the single pollination experiment. Mixed effect models were applied where required, otherwise two-way ANOVA and TukeyHSD *post hoc* test were used. Upper-case letters (above the line) indicate statistical differences dependent on the maternal genotype, lower-case letters (below the line) indicate statistical differences dependent on the pollen donor, *P* < 0.05. (d-f) Spearman’s correlation analysis between seed size and seed mass (d), seed size and seed number (e) and seed number and (f) seed mass. The respective correlation coefficients (*R*) and p-values (*P*) are displayed on each graph.

## Results

### The magnitude of PPEB only partially corresponds with the expression of SLRs

To verify whether pollination with favored and non-favored pollen results in predictable changes in the magnitude of PPEB and to determine if this variation is dependent on the SLR expression in the parental genotypes, we measured the ethylene emission after hand pollination of different maternal accessions with pollen donors selected from each SLR expression group as described in the ‘*Single pollination experiment*’ section. Natural accession G2 and the transgenic line *irSLR1*/Ut have both lost the ability of mate selection due to the lack of the stylar expression of SLR1 and SLR2. As expected, no significant differences in PPEB were observed after pollination of those maternal genotypes with the selected pollen donors (Fig 1c). Pollination of EV-SOL (SLR1 expression) with line 274 (no SLR expression) resulted in significantly lower PPEB compared to the other two pollen donors. As EV-SOL has been shown to select strongly against the accessions with no SLR expression (Guo *et al*., 2019), this PPEB phenotype corresponds with the mate selection pattern observed for this line. Pollination of the transgenic line silenced in the expression of SLR2 (irSLR2/Az) with pollen donor 274 resulted in a PPEB comparable to the one observed after pollination on this line with pollen donor 194 that expresses only SLR1, but it was significantly higher after pollination with line 305 expressing both SLRs, a phenotype identical with the one observed for AzWT (Fig. 1c) which has been shown to select strongly for the pollen donors with similar SLR expression pattern (Guo *et al*., 2019). Therefore, the higher PPEB after pollination of this genotype with line 305 corresponds to AzWTs preference for pollen donors expressing both SLRs which is not the case for line *irSLR2*/Az. As *irSLR2*/Az shows a PPEB phenotype similar to that of AzWT despite expressing only SLR1, like *EV-SOL*, we conclude that the control of PPEB only partially correlates with the expression of SLRs and is more complex than initially suspected. The fact that both *irSLR2*/Az and AzWT produced, on average, more ethylene than the other three maternal genotypes despite having different SLR expression profiles, indicates that additional elements must be involved in the regulation of PPEB.

### High level of PPEB correlates with the lack of SLR2 expression

To investigate if and how the PPEB influences the transcript and protein abundance of the two SLRs we quantified the PPEB in 650 recombinant inbred lines (RILs) from the MAGIC population after hand pollination with the pollen from Utah wild type accession (Fig. S1). Subsequently, we selected a subset of individuals with extreme phenotypes for further testing. After reconfirming the magnitude of PPEB in these lines (Fig. 2a), we measured the stylar expression and protein abundance of both SLR1 (Fig. 2b-d) and SLR2 (Fig. 2f-g). SLR1 expression and the protein were detected in nine lines with the low PPEB and nine lines with the high PPEB. However, gene expression levels and protein abundance of SLR2 were detected in five lines with low PPEB and only one high PPEB line. We computed the Spearman’s rank correlation coefficient to assess the relationship between PPEB and the SLRs (Table S2). There was no significant correlation for SLR1, but we observed a significant negative correlation between PPEB and the expression and protein abundance of SLR2 (*R* = -0.38, *R* = -0.38, respectively, *P* < 0.05). These results illustrate that a high level of PPEB after pollination with a standard pollen donor is associated with the lack of stylar expression of SLR2. As not all of the tested lines followed this pattern, these findings further support the involvement of additional regulatory mechanisms in the control of mate selection.

### Variation in the tradeoff between offspring size vs. number underlines the differences in reproductive strategies within the species

As *N. attenuata* flowers consistently select mates with corresponding SLR expression pattern among genotypes of non-self pollen, we hypothesized that this mechanism may have evolved to select particular mates with beneficial traits linked to certain SLR expression patterns. To test this hypothesis, we pollinated the three previously selected maternal genotypes with the pollen donor representatives of the three SLR expression groups (see the ‘*Single pollination experiment*’ section) and examined different seed attributes of the offspring (Fig. 3). No significant differences associated with the pollen donor were observed for seed mass and seed number. Among the three maternal genotypes, EV-SOL generally showed the lowest seed mass and seed number while G2 showed the highest (Fig. 3a,b). The strong positive correlation between those two traits was confirmed by a Spearman’s rank correlation coefficient (R = 0.92, *P* < 2.2e – 16; Fig. 3f). Average seed size correlated negatively with seed mass (R = -0.46, *P* = 6.1e – 06; Fig. 3d) and seed number (R = -0.71, *P* = 1e – 14; Fig. 3e), indicating a tradeoff between offspring size and number. Indeed, the low seed count for EV-SOL resulted in larger seeds, while the high seed count in G2 resulted in the smallest seeds among the three genotypes tested. Even though AzWT produced significantly more seeds than EV-SOL, they were of a similar size (Fig. 3b,c). G2 was the only maternal genotype that showed significant differences in the average seed size as a function of the pollen donor, with the largest seeds produced in the cross with line 274 and the smallest seeds produced in the cross with line 305 (Fig. 3c). Together, these results illustrate that within the same species different maternal genotypes may express different reproductive strategies.

### Mate selection mechanisms affect the germination time of the progeny

To further investigate the effect of mate choice on the offspring, we examined the emergence and early seedling establishment up to 14 days after sowing (DAS). In this experiment, we also included the self-pollinated seeds as an additional reference. The fastest germinating seeds were produced by G2 (Fig. 4). Within three DAS, G2 showed 100% germination in the cross with line 274, 97% when self-pollinated, and 90% when crossed with line 194. The seeds obtained from the cross of G2 with line 305 germinated marginally more slowly, with fewer than half of the seeds germinating within three DAS. Four DAS, 97% of the seeds from this cross were germinated. For AzWT 97% of seeds germinated within three DAS in the cross with line 194, 90% of seeds germinated in the cross with line 274 and only 77% of the seeds germinated in the cross with line 305. The self-pollinated seeds of AzWT showed the slowest germination, with most of the seedlings emerging four DAS. In contrast, the seeds produced by EV-SOL germinated much more slowly, especially for the seeds obtained after self-pollination, where only a few seedlings emerged four DAS and only 83% of them were germinated six DAS. The best-performing offspring of this maternal genotype were produced in the cross with the non-favored line 274, where 87% of seeds had already germinated four DAS.

**Fig. 4:**
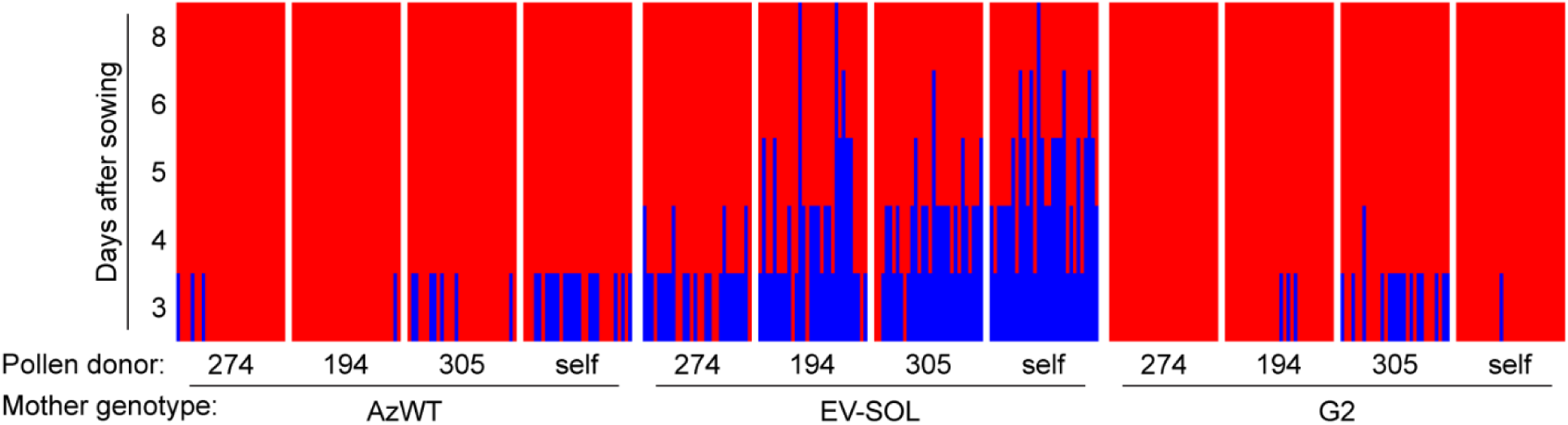
In *Nicotiana attenuata* the germination time depends on maternal genotype and the pollen donors. A heatmap showing the germination time of 30 seeds from three different capsules per plant produced by maternal genotypes after single hand pollination with a favored, non-favored or neutral pollen donor. The germinated seeds are shown in red and the non-germinated seeds are shown in blue. Germination time was recorded for each seed daily, starting three days after sowing until the majority of seeds had germinated.

### Seedling establishment and early growth dynamics depend on both parental genotypes

To evaluate if the variation in seed size and number as well as in the germination time affects the seedling establishment, we measured the shoot- and root-related traits of the seedlings from the different crosses 14 DAS (Figs 5, S2). Additionally, we monitored the growth of the seedlings every two days to provide a detailed description of the early growth dynamics (Figs S3, S4). The best performing offspring in this experiment was obtained from a cross of EV-SOL and line 274. The seedlings obtained from this cross exceeded in all the traits measured in this experiment except for the hypocotyl length. They showed the longest tap root and the highest number of lateral roots (Fig. S4b), resulting in the highest root biomass. These individuals entered the logarithmic phase of growth faster than all the other seedlings (Fig. S3) and produced the largest rosettes and the highest shoot biomass. On the other hand, the seedlings obtained from the cross between AzWT and its favored pollen donor (line 305) were consistently slightly bigger than those from the other pollen donors (Figs 5, S2-S4). However, since only the results observed for the root length were statistically significant, this tendency would have to be further verified. The cross of G2 with line 194 produced fewer, but longer lateral roots, whereas the cross with line 305 resulted in a large number of short lateral roots demonstrating, that differences in early root architecture depend on the pollen donor (Figs 5, S4b). G2 seedlings were also characterized by very long hypocotyls, especially after self-pollination (Fig. 5d). Taken together, these results indicate that the mate selection mechanisms and different reproductive strategies observed in *N. attenuata* result in differences in offspring performance that are visible already at the early stages of seedling establishment.

**Fig. 5:**
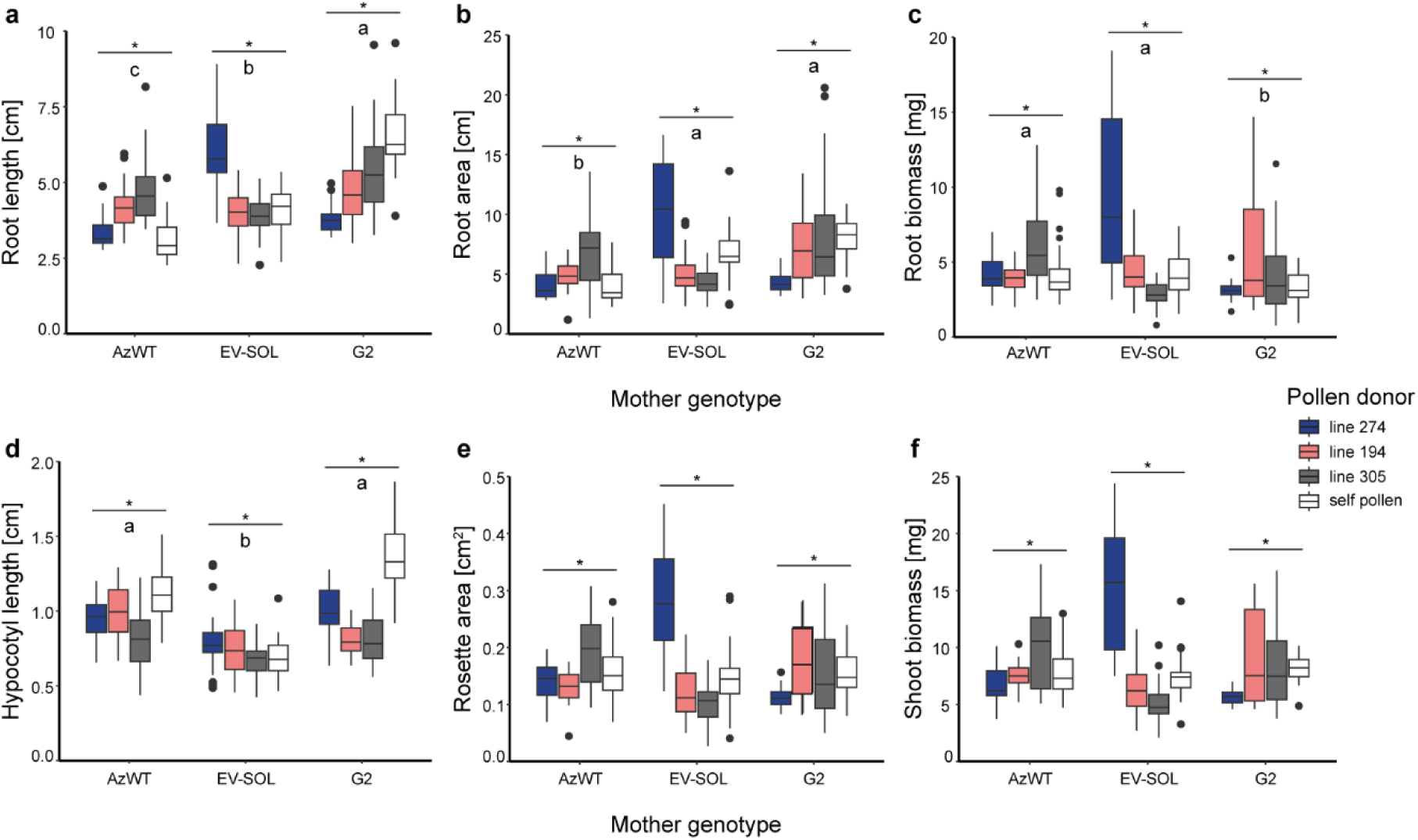
Early establishment of *Nicotiana attenuata* seedlings depends on maternal genotype and the pollen donors. (a-c) Root- and (d-f) shoot-related traits were determined for the offspring of different maternal genotypes after single hand pollination with a favored, non-favored, neutral or self pollen donor, 14 days after sowing (DAS). (a) Root length (the length of the tap root), (b) root area (the sum of lengths of tap root and secondary roots), (d) hypocotyl length, and (e) rosette area were measured from scans of the seedlings using Fiji software. (c) Root and (f) shoot biomass were measured destructively at the end of the experiment. Data are shown as means of up to 30 seedlings per cross - ten randomly selected seeds from each of the different capsules obtained from single pollination experiment (related data in Figs 3, 4). Statistical differences were calculated using mixed effect models with a Sidak *post hoc* test to extract multiple comparisons; small letters (below the line) indicate statistical differences dependent on the pollen donor, *P* < 0.05.

### GWAS identifies a major peak associated with variation in PPEB

To explore the genetic architecture underlying mate selection and identify novel genes involved in its regulation, we conducted a GWAS on PPEB data obtained from the 26-parent *N. attenuata* MAGIC population. The quantification of the PPEB after reciprocal single hand pollination of nearly 650 RIL lines (see the ‘*Genome Wide Association Study (GWAS)*’ section) resulted in two distinct datasets (Fig. S1): one with varied maternal genotypes and standardized pollen donor that we used to impute style-expressed genes and to elucidate the role of the maternal genotype in the mate selection processes and one in which a standardized maternal genotype and varied pollen donors was used to impute pollen-related genes and explore signals derived from different pollen donors. We found no significant association from the analysis of the dataset with the standardized maternal genotype, whereas the GWAS of the dataset with varied maternal genotypes resulted in the identification of nine significant associations located on chromosome 9 and one association located on chromosome 4 (Fig. S5, Table S4). The phenotypic variance explained by the individual significant SNPs was c. 10%. Among the nine significant SNPs identified on chromosome 9, eight were part of a larger, sharply defined and clearly identified peak of associations covering c. 14.5 Mbp on chromosome 9 (Fig. S5a). Therefore, in our search for the candidate genes, we included additional markers that were part of this peak, though slightly below the significance threshold (Table S4). Assuming a linkage disequilibrium (LD) decay distance of 2.5 Mbp on either side of each significant marker identified a large number of genes. The list of potential candidates was narrowed down to the genes specifically expressed in the female reproductive organs based on *N. attenuata* eFP browser (Brockmöller *et al*., 2017). Among those, seven final candidates including an ethylene-responsive transcription factor (*ERF061*), two putative leucine-rich receptor-like protein kinases, an *N. attenuata* homolog to an early nodulin-like proteins previously identified in *A. thaliana,* and a cluster of three genes annotated as regulators of nonsense transcripts 1 were selected based on functional annotations and the current state of the literature (Table S5). We propose that these seven candidates are likely to be involved in the regulation of mate selection in *N. attenuata* and should be analyzed in more detail.

## Discussion

Disruption of ethylene signaling in ethylene-deficient mutant lines and the lack or loss of the expression of the *Na*S-like RNases (SLRs), have both been shown to result in the loss of mate selection abilities (Bhattacharya & Baldwin, 2012; Guo *et al*., 2019), but have not previously been investigated for potential links in their function. Therefore, we first explored if and how ethylene signaling affects the SLR expression and/or activity. We found that the PPEB is only partially correlated with the SLR expression and that some of the observed variation depends on the genetic background of the maternal genotype rather than the SLR expression pattern (Fig. 1c). Moreover, our results showed a correlation between high PPEB and the lack of the expression of SLR2 (Fig. 2, Table S2). The lack of any specific correlation between the magnitude of PPEB and the transcript and protein abundance of SLR1 points to a difference in the regulation of the two SLRs and possibly in their function in facilitating the pre-zygotic mate selection. It has been shown that variation in the SLR protein abundance among natural *N. attenuata* accessions is correlated with DNA methylation (Guo *et al*., 2019). Additionally, a recent study suggested that in the absence of a sufficient amount of ethylene, the growth of the pollen tube might be blocked through a mechanism involving the ethylene receptors (ETRs), leading to the modification of pollen tube cell walls and Ca^2+^ loading (Althiab-Almasaud *et al*., 2021). A calcium-dependent protein kinase that phosphorylates S-RNase (Nak-1) has been identified in *Nicotiana alata*, but its specific function in SI is not clear (Kunz *et al*., 1996). Differential phosphorylation/dephosphorylation of the two SLRs in addition to the variation in the SLR protein abundance in natural accessions could explain the variation observed in our study and should be subjected to further research.

To address the longstanding question of whether the mate choice mechanisms are adaptive, resulting in a selection of mates with beneficial traits and ultimately leading to the production of superior progeny, we examined the performance of the offspring obtained after pollination with favored and non-favored pollen donors. The analysis of the differences in the seed size and seed number per capsule among the different crosses revealed the existence of a tradeoff between offspring size and number (Fig. 3). Such offspring size vs. number tradeoffs are common among the plant and animal kingdoms (Fox & Czesak, 2000; Gnan *et al*., 2014; Dani & Kodandaramaiah, 2017) and yet our results show that their level may vary between different genotypes, highlighting the differences in reproductive strategies within the same species. Our results suggest that the *N. attenuata* accessions that are able to select for the favored pollen show a consistent tendency to produce larger and faster growing progeny after the pollination with the favored pollen donors (Fig. 5). In contrast, the selection for beneficial traits is clearly abolished when the maternal genotype is only able to discriminate against certain pollen donors, e.g., in the genotype expressing only SLR1. In this case, the best-performing progeny produced by this maternal genotype and the best-performing progeny overall was obtained from the cross with the non-favored pollen donor (Figs 5, S2). The pollen donors that do not express any SLRs have been shown to be strongly selected against by the maternal genotypes that express the SLRs, regardless of the exact expression pattern (Guo *et al*., 2019). Therefore, we speculate that the rejection of pollen tubes from these donors could be correlated with the expression of SLR1 specifically, further supporting our hypothesis that the two SLRs differ in their function in mate selection, e.g., SLR1 could inhibit the growth of pollen tubes while SLR2 could promote it. To test this hypothesis, pollen tube growth rates would have to be measured in different crosses including the genotypes that express only SLR2, which were not included in this study due to lack of existing knowledge on their mate selection patterns. Alternatively, it is possible that the loss of SLRs was accompanied by the loss of other elements linked to the S-locus that are necessary for the mate selection. As a consequence, the pollen tubes from those accessions would be inhibited by the maternal genotypes that express SLRs due to the lack of necessary interacting proteins delivered by the pollen, regardless of the potential benefit of producing superior offspring. The exact molecular mechanism of mate selection is still not known, but an obvious first step to test the hypothesis would be to investigate the expression of pollen-specific S-determinants as well as genes homologous to non-S-locus modifiers and their interaction with the two SLRs in the parental genotypes. As the reproductive success of an individual is measured not only by the number of their offspring, but also by their offspring’s ability to produce further offspring of their own more research is needed to determine whether mate selection is adaptive.

In the experiment described here, the non-favored pollen produced the best-performing offspring which supports the hypothesis that the SI elements are being pseudogenized after the SI-SC transition of *N. attenuata,* resulting in the reduced fitness of the accessions expressing SLRs (Guo *et al*., 2019). This hypothesis is also supported by the results observed for the offspring of the non-selecting maternal line, G2 (Fig. 5), for which the offspring size vs. number tradeoff was much more pronounced, but its direction changed compared to the other two maternal genotypes (Fig. 3). We also observed significant differences in seed size between different pollen donors, suggesting differential maternal investment in the genetically variable offspring and possibly a higher level of parent-offspring conflict (Smith & Fretwell, 1974). This hypothesis requires further testing, perhaps by comparing the seed metabolome after single and mixed pollinations or through the analysis of maternal and zygotic genetic controls on seed size and number (Van Daele *et al*., 2012). As a result of adopting a different reproductive strategy, the non-selecting line produced a large number of small seeds with different genetic qualities, contributing to better overall performance of its progeny. Based on the phenotypic traits and early growth dynamics observed for the seedlings of this maternal genotype, we concluded that they were more efficient in transitioning from the heterotrophic to autotrophic lifestyle (Zacchello *et al*., 2020) and better equipped to compensate for limiting resources in the environment and aboveground competition (Mašková & Herben, 2018). This kind of phenotypic plasticity among the offspring at the early stages of development would be beneficial in the context of the natural history of *N. attenuata*. The fire chasing germination behavior of this species results in long dormancy periods and subsequent germination and growth in genetically diverse populations (Bahulikar *et al*., 2004). Hence, producing a large number of genetically diverse offspring might amount to a potential bet-hedging strategy, with implications for the survival of long-lived seed banks and accounting for different growth conditions after germination (Anderson *et al*., 2023).

The results described here suggest that the choice of mates observed in mixed, non-self pollination experiments leads to the production of offspring with different growth attributes. We present evidence that even though the PPEB correlates to a certain degree with the SLR expression pattern in the parental genotypes, it is likely that additional elements and regulatory mechanisms are involved in the control of this process. Among the significant associations obtained from the pollination of different maternal genotypes with a standardized pollen donor, we identified seven potential candidates (Table S5). The gene encoding the ethylene-responsive transcription factor (*ERF061*, Niat3g_65650) is particularly interesting because of its strong expression in the style and stigma, whereas an *A. thaliana* homolog of this gene is expressed in the papillae cells and siliques. No functional analysis of this gene has been reported to date, but its highly specific expression profile indicates its potential involvement in the early stages of pollen-pistil interaction, which is where pre-zygotic mate selection occurs (Bhattacharya & Baldwin, 2012). Similarly, LRR protein kinases have been recently shown to be involved in early stages of pollen recognition and pollen-pistil interactions in *A. thaliana* (Lee & Goring, 2021). The involvement of receptor kinases in the later stages of the reproduction, such as pollen tube guidance to the ovule, or its reception, have been already described in great detail (Zhong & Qu, 2019; Zhou & Dresselhaus, 2019), but the research has mainly focused on the model system *A. thaliana.* Further characterization of the LRR protein kinases identified in our analysis (Niat3g_66018 and Niat3g_65722) could provide novel insight into the function of those proteins in sexual reproduction of different families. In contrast, the gene annotated as the early nodulin-like protein 3 (*ENOD3*, Niat3g_65615) is expressed specifically in ovaries and flower buds, suggesting that it may act at later stages of male-female communication. Indeed, a group of female-specific early nodulin-like proteins have been identified in *A. thaliana*, where they have been shown to be involved in the pollen tube entrance into the ovule, growth arrest and burst (Hou *et al*., 2016). Finally, we identified a cluster of three genes annotated as regulators of nonsense transcript 1 (Niat3g_65293, Niat3g_65295 and Niat3g_65296). These genes share a high homology with UPF1, an RNA helicase involved in the non-sense mutation mediated RNA decay (NMD). In *A. thaliana*, UPF1 has been shown to act as a maternal factor controlling seed size (Yoine *et al*., 2006). The NMD factors such as UPF1 and SMG7 are involved in the regulation of a number of physiological processes in plants, including ethylene signaling, meiosis, seed set and seed size control (Raxwal *et al*., 2020). There is growing evidence for a wide range of non-canonical functions of these proteins that expand beyond RNA quality control (Raxwal & Riha, 2023). Besides the potential role of NMD in supporting the degradation of RNA of the incompatible pollen, UPF1 or NMD could be involved in fine-tuning the expression of other genes related to mate selection or act via ethylene signaling. Most of the studies on NMD in plants are based on *A. thaliana* and *Physcomitrella patens.* Unravelling the putative role of NMD processes in ethylene signaling and mate selection in *N. attenuata* will allow a better understanding of molecular mechanisms controlled by non-canonical functions of NMD factors and plant sexual reproduction.

## Supporting information

Baraniecka_et_al_2023_supporting_information

## Acknowledgments

We thank Prof. Sarah E. O’Connor for supporting the project, Evelyn Claußen, Mohammed Abdulazeez and Melanie Smith for technical support, Dr. Grit Kunert for statistical advice, Dr. Rayko Halitschke for help with ethylene measurements and scientific support, the glasshouse team for the plant cultivation and the members of the department for their help with germination and harvesting of the MAGIC population. This work was supported by the Max Planck Society and the Collaborative Research Centre “Chemical Mediators in Complex Biosystems-ChemBioSys” (SFB 1127) to ITB.

## Author contributions

PB, ITB and JRP provided the intellectual framework for the study, PB and WS designed and performed the experiments with the help of EM, PB and EM analyzed the data. The original manuscript was prepared by PB with the help of KG, DK, WS and KG. All authors contributed intellectual input to this study as well as reviewed and agreed to this manuscript.

