## Supplementary material for "Pre-zygotic mate selection affects progeny fitness and is only partially correlated with the expression of *Na*S-like RNases": Baraniecka_et_al_2023_supporting_information

The following Supporting Information is available for this article:

**Fig. S1** Quantification of the post-pollination ethylene burst (PPEB) in 650 recombinant inbred lines (RILs) from a 26-parent Multiparent Advanced Generation Inter-Cross (MAGIC) population of *Nicotiana attenuata*.

**Fig. S2** Variation in *Nicotiana attenuata* seedling phenotype of different parental genotype combinations.

**Fig. S3** Early growth dynamics of *Nicotiana attenuata* seedlings.

**Fig. S4** Number of leaves and secondary roots of *Nicotiana attenuata* seedlings 14 days after sowing (DAS).

**Fig. S5** Manhattan plots summarizing the results of the Genome Wide Association Study (GWAS) on the post-pollination ethylene burst (PPEB) in *Nicotiana attenuata*.

**Table S1** List of primers used in this study

**Table S2** Spearman's rank correlation coefficients between the post-pollination ethylene burst (PPEB) and the gene expression and protein abundance of SLR1 and SLR2 in *Nicotiana attenuata*.

**Table S3** Genome Wide Association Study (GWAS) results for the SNPs associated with the post-pollination ethylene burst (PPEB) in the *Nicotiana attenuata* 26-parent Multiparent Advanced Generation Inter-Cross (MAGIC) population after pollination with Utah wild type (UtWT) standard pollen donor.

**Table S4** Genes selected as candidates for further research after the Genome Wide Association Study (GWAS) on post-pollination ethylene burst (PPEB) in *Nicotiana attenuata*.

**Fig. S1** Quantification of the post-pollination ethylene burst (PPEB) in 650 recombinant inbred lines (RILs) from 26-parent Multiparent Advanced Generation Inter-Cross (MAGIC) population of *Nicotiana attenuata*. The PPEB data obtained after single hand pollination of each RIL line with the pollen from Utah wild type accession are shown in grey. Lines selected for further analyses of the correlation between ethylene emission and SLR expression and protein abundance (Fig. 2) are highlighted: lines with the lowest PPEB in blue and lines with the highest PPEB in red. The PPEB varied in range from 37.78 to 801.75 ppbv. The PPEB data obtained after single hand pollination of Utah wild type with the pollen from each RIL line are shown in green. The PPEB varied in range from 22.89 to 881.26 ppbv. Separate ordering of the RILs (x-axis) was performed for each dataset in order to plot PPEB (y-axis).

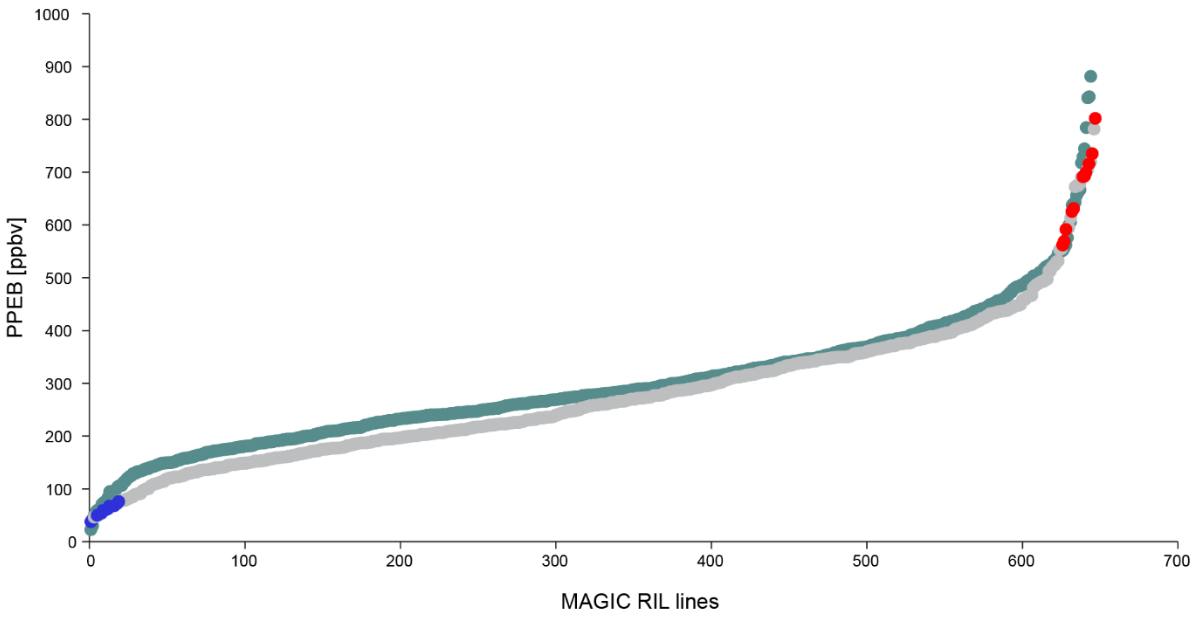

**Fig. S2** Variation in *Nicotiana attenuata* seedling phenotype of different parental genotype combinations. Representative seedlings of three different maternal genotypes after the pollination with a favored, non-favored or neutral pollen donor photographed 14 days after sowing, before the harvest.

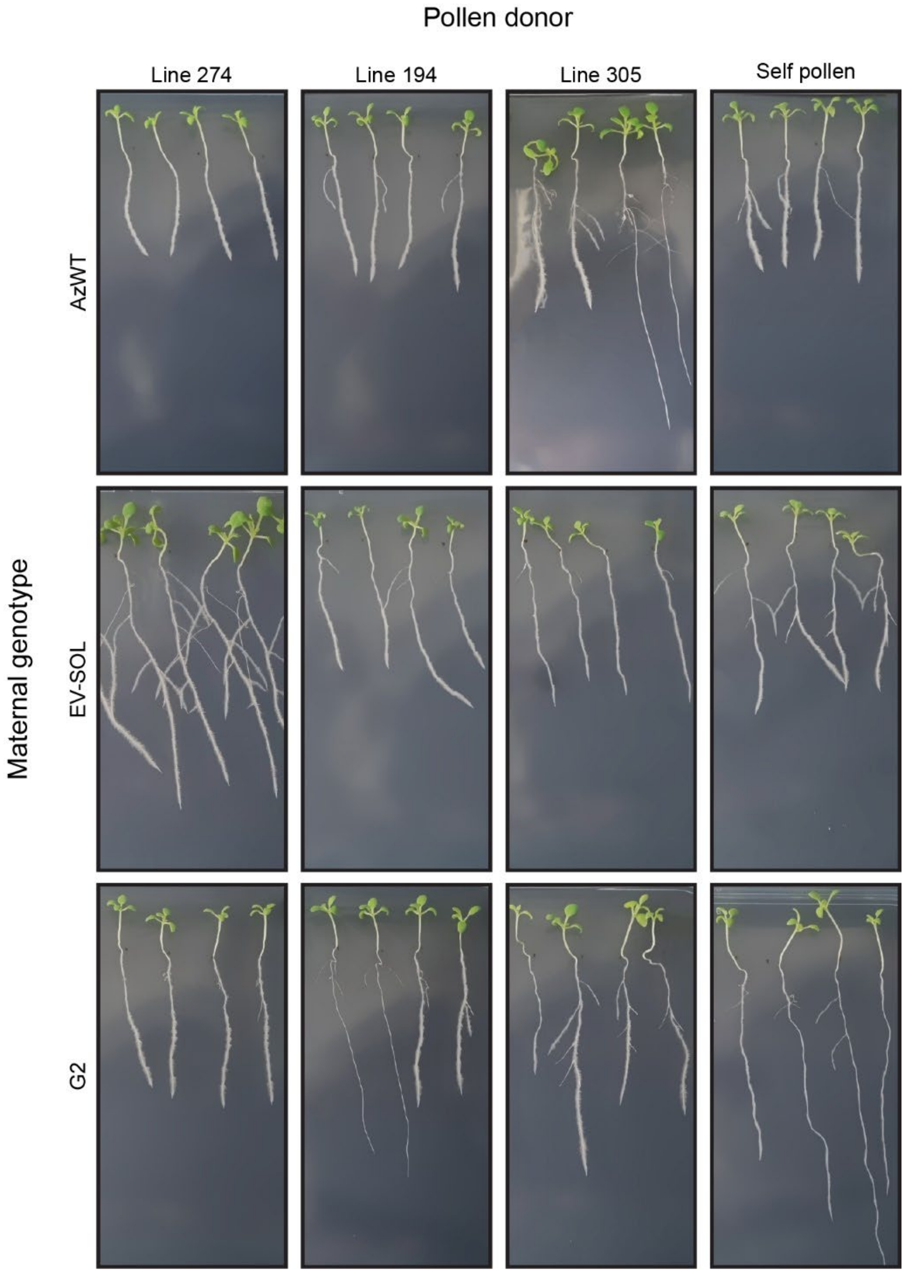

**Fig. S3** Early growth dynamics of *Nicotiana attenuata* seedlings. The early growth dynamics were determined for the offspring of different maternal genotypes after single hand pollination with a favored, non-favored, neutral or self pollen donor. (a) Root length (the length of tap root), (b) root area (the sum of lengths of tap root and secondary roots), (c) hypocotyl length, and (d) rosette area were measured from scans of the seedlings using ImageJ. The seedlings were scanned every second day starting 4 days after sowing (DAS) for 14 days. Data are shown as means of up to 30 seedlings per cross - 10 randomly selected seeds from the three different capsules obtained from the single pollination experiment (related data in Figs 3-5). Statistical differences were calculated 14 DAS using mixed effect models with a Sidak *post hoc* test to extract multiple comparisons; small letters (below the line) indicate statistical differences dependent on the pollen donor, $P <$ 0.05.

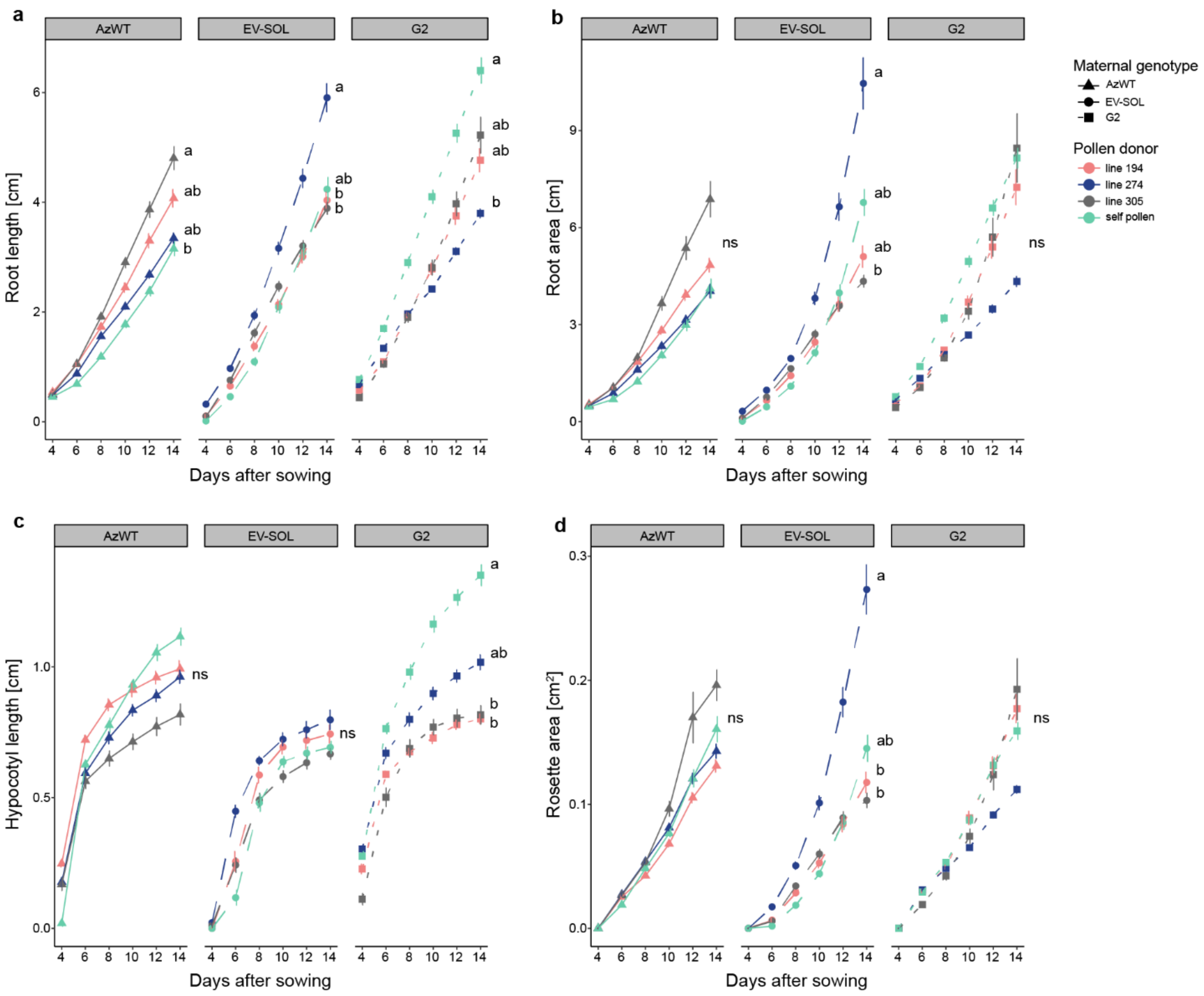

**Fig. S4** Number of leaves and secondary roots of *Nicotiana attenuata* seedlings 14 days after sowing (DAS). (a) Number of leaves and (b) lateral roots were determined for the offspring of different maternal genotypes after single hand pollination with a favored, non-favored, neutral or self pollen donor. The seedlings were scanned every second day starting 4 DAS for 14 days and the scans were used to extract the data.

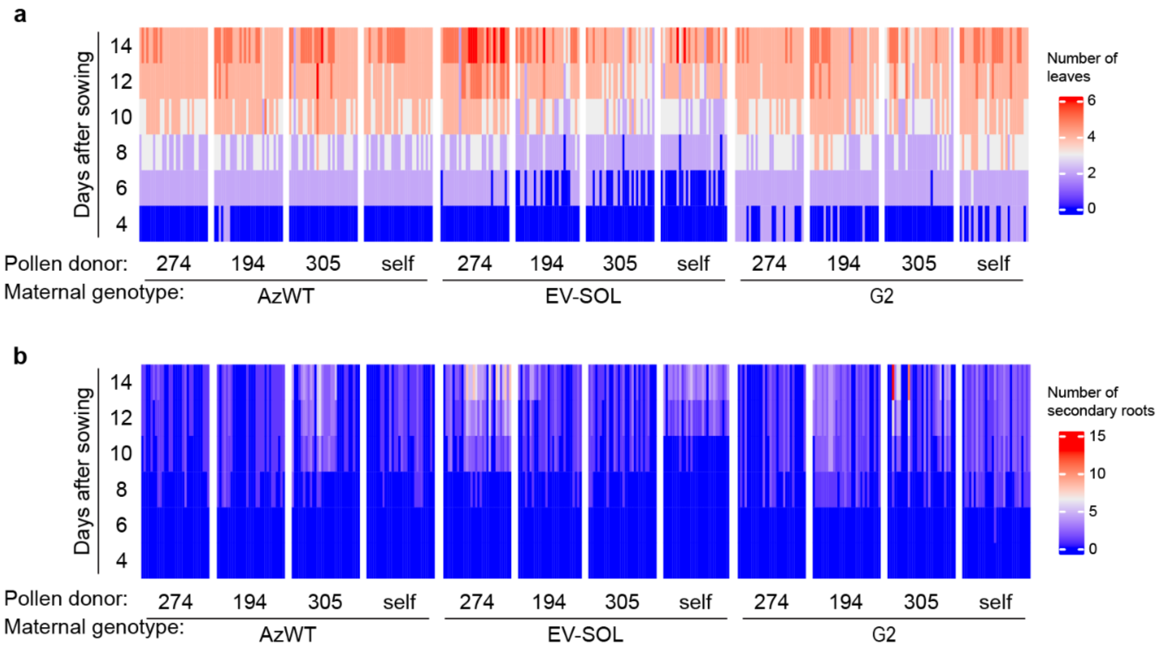

**Fig. S5** Manhattan plots summarizing the results of the Genome Wide Association Study (GWAS) on the post-pollination ethylene burst (PPEB) in *Nicotiana attenuata*. The results of the GWAS on PPEB obtained after reciprocal the pollination of (a) different maternal genotypes (629 RIL lines from the MAGIC population) with a Utah wild type (UtWT) standard pollen donor and (b) the standard maternal genotype (UtWT) with different pollen donors (629 RIL lines from the MAGIC population). The strongest association peak on chromosome 9 is shown in blue. Additional significantly associated markers are shown in red. The horizontal dashed line indicates the significance threshold set to 1e-05. The x-axis indicates the position of each SNP in the genome and the y-axis shows the negative logarithm of the p-value obtained from the GWAS model. Different linkage groups are shown in different colors. Scf – scaffold of the genome assembly.

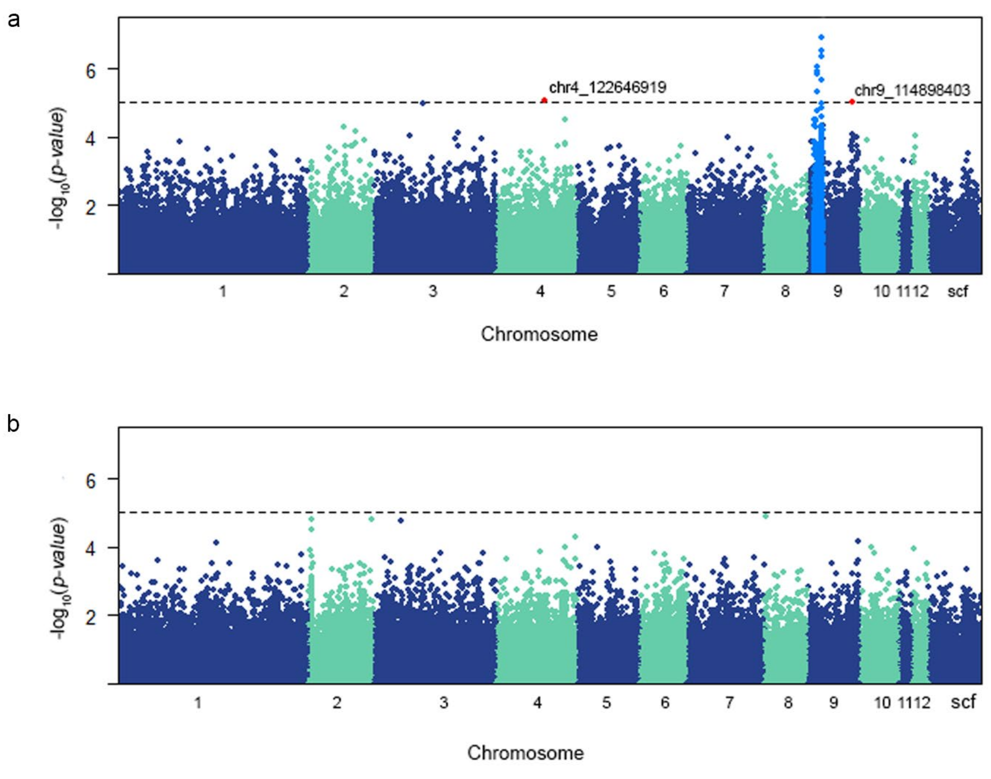

**Table S1** List of primers used in this study

| Gene | Primer name | Sequence (5' to 3') |
|---|---|---|
| *Na* SLR1 | SLR1-RT-F | CACATTTAACGAAACCAGAGATGCC |
| *Na* SLR1 | SLR1-RT-R | TTTATATGTTCGACGCACTTGAGGT |
| *Na* SLR2 | SLR2-RT-F | AGCGTATCCTAACCTCAATTGCATT |
| *Na* SLR2 | SLR2-RT-R | GTTGCCTCTCGGTTGAAACAGA |
| *Na* EF1a | EF1a-F | CCACACCTCCCACATTGCTGTCA |
| *Na* EF1a | EF1a-R | CGCATGTCCCTCACAGCAAAAC |

**Table S2** Spearman's rank correlation coefficients between the post-pollination ethylene burst (PPEB) and the gene expression and protein abundance of SLR1 and SLR2 in *Nicotiana attenuata*. Significant correlations are underlined.

| | PPEB : SLR1 | | PPEB : SLR2 | |
|---|---|---|---|---|
| | *R* | *P* | *R* | *P* |
| Relative expression | 0.13 | 0.19 | **<u>-0.38</u>** | **<u>4.1e-05</u>** |
| Protein abundance | -0.026 | 0.79 | **<u>-0.38</u>** | **<u>6.6e-05</u>** |

**Table S3** Genome Wide Association Study (GWAS) results for the SNPs associated with post-pollination ethylene burst (PPEB) in the *Nicotiana attenuata* 26-parent Multiparent Advanced Generation Inter-Cross (MAGIC) population after pollination with Utah wild type (UtWT) standard pollen donor. The table provides the SNP ID (SNP), linkage group (Chromosome), base pair position of the marker (Position), p-value (*P*), minor allele frequency (MAF), the amount of variation explained by each marker ($R^2$), p-value following the procedure controlling the false discovery rate (FDR) and the allelic effect estimate (Allelic effect). Sample size = 629. The significant associations were declared based on an LOD threshold of 1e-05 and are shown above the dashed line. Additional SNPs associated with the peak on chromosome 9 which were used for the selection of candidate genes are shown below the dashed line.

| SNP | Chromosome | Position | *P* | MAF | $R^2$ | FDR | Allelic effect |
|---|---|---|---|---|---|---|---|
| chr9_33910631 | 9 | 33910631 | 1.19E-07 | 0.334658188 | 0.109138598 | 0.021865296 | 46.28255771 |
| chr9_34550003 | 9 | 34550003 | 3.15E-07 | 0.453895072 | 0.106077878 | 0.028853351 | 39.9471141 |
| chr9_33911093 | 9 | 33911093 | 4.71E-07 | 0.314785374 | 0.104815634 | 0.028853351 | 42.85446266 |
| chr9_23566194 | 9 | 23566194 | 8.65E-07 | 0.322734499 | 0.102913666 | 0.039778506 | 42.66627007 |
| chr9_23566261 | 9 | 23566261 | 1.16E-06 | 0.321144674 | 0.102003536 | 0.042618002 | 42.27956624 |
| chr9_24668222 | 9 | 24668222 | 1.46E-06 | 0.355325914 | 0.101280458 | 0.044808013 | 38.93679848 |
| chr9_34639866 | 9 | 34639866 | 2.08E-06 | 0.466613672 | 0.100180575 | 0.054731257 | -36.30640923 |
| chr9_24958799 | 9 | 24958799 | 4.83E-06 | 0.461844197 | 0.097578978 | 0.111043997 | -31.42123162 |
| chr4_122646919 | 4 | 122646919 | 8.39E-06 | 0.320349762 | 0.095877018 | 0.160358583 | 40.39846815 |
| chr9_114898403 | 9 | 114898403 | 9.90E-06 | 0.235294118 | 0.095369509 | 0.160358583 | -38.09307256 |
| chr9_34540947 | 9 | 34540947 | 1.04E-05 | 0.470588235 | 0.095231257 | 0.160358583 | 33.78920023 |
| chr9_34527745 | 9 | 34527745 | 1.44E-05 | 0.471383148 | 0.094228425 | 0.203311507 | 33.37371145 |
| chr9_22742053 | 9 | 22742053 | 1.75E-05 | 0.368044515 | 0.093627995 | 0.229738193 | -38.32508298 |
| chr9_34538085 | 9 | 34538085 | 2.59E-05 | 0.46581876 | 0.092427596 | 0.300710507 | 30.58005968 |
| chr9_34641035 | 9 | 34641035 | 2.62E-05 | 0.466613672 | 0.092399026 | 0.300710507 | 31.71154558 |
| chr9_15468213 | 9 | 15468213 | 3.07E-05 | 0.321144674 | 0.091915527 | 0.312560159 | 37.48807796 |
| chr9_20664570 | 9 | 20664570 | 3.08E-05 | 0.417329094 | 0.091901099 | 0.312560159 | -33.03842207 |
| chr9_20251720 | 9 | 20251720 | 3.40E-05 | 0.353736089 | 0.091602273 | 0.312560159 | 34.78062146 |
| chr9_37752304 | 9 | 37752304 | 4.66E-05 | 0.390302067 | 0.090646102 | 0.36831735 | 32.08830099 |
| chr9_33910755 | 9 | 33910755 | 4.66E-05 | 0.352941176 | 0.090643397 | 0.36831735 | 33.10267059 |
| chr9_14312849 | 9 | 14312849 | 4.74E-05 | 0.309220986 | 0.090591666 | 0.36831735 | 36.5187105 |
| chr9_34550760 | 9 | 34550760 | 4.97E-05 | 0.475357711 | 0.090448076 | 0.36831735 | 29.75647313 |
| chr9_21530138 | 9 | 21530138 | 5.27E-05 | 0.461844197 | 0.09026941 | 0.373028753 | 31.34252268 |
| chr9_34552619 | 9 | 34552619 | 5.74E-05 | 0.476947536 | 0.090012474 | 0.385315009 | 29.51279708 |
| chr9_33910882 | 9 | 33910882 | 5.87E-05 | 0.321144674 | 0.089946942 | 0.385315009 | 32.23052847 |
| chr9_34530934 | 9 | 34530934 | 6.42E-05 | 0.468203498 | 0.089674598 | 0.398213169 | 28.88884572 |
| chr9_36956308 | 9 | 36956308 | 6.49E-05 | 0.388712242 | 0.089638667 | 0.398213169 | 32.0423806 |
| chr9_34615976 | 9 | 34615976 | 7.51E-05 | 0.457074722 | 0.089198218 | 0.431914842 | 29.11170309 |
| chr9_34945282 | 9 | 34945282 | 8.66E-05 | 0.362480127 | 0.088769985 | 0.439198259 | 33.18234032 |
| chr9_34648379 | 9 | 34648379 | 8.72E-05 | 0.472972973 | 0.088747846 | 0.439198259 | 29.61976025 |

**Table S4** Genes selected as candidates for further research after the Genome Wide Association Study (GWAS) on post-pollination ethylene burst (PPEB) in *Nicotiana attenuata*. The table provides the gene code (GOI), gene name and annotation, the position of the gene on chromosome 9, the most significantly associated marker and its p-value. LOD threshold = 1e-05.

| GOI | Name | Annotation | Position | | Marker | *P* |
|---|---|---|---|---|---|---|
| | | | Start | End | | |
| Niat3g_65650 | ERF061 | Ethylene-responsive transcription factor 061 | 25738069 | 25738948 | chr9_23566194 | 8.65E-07 |
| Niat3g_66018 | LRR1 | Putative LRR protein kinase | 35002162 | 35006145 | chr9_33910631 | 1.19E-07 |
| Niat3g_65722 | LRR2 | Putative LRR protein kinase | 26738712 | 26742144 | chr9_24668222 | 1.46E-06 |
| Niat3g_65615 | ENL3 | Early nodulin-like protein 3 | 24287027 | 24288294 | chr9_23566194 | 8.65E-07 |
| Niat3g_65293 | RNT1a | Regulator of nonsense transcripts 1-like protein | 17322262 | 17324573 | chr9_15468213 | 3.07E-05 |
| Niat3g_65295 | RNT1b | Regulator of nonsense transcripts 1-like protein | 17326468 | 17329564 | chr9_15468213 | 3.07E-05 |
| NIat3g_65296 | RNT1c | Regulator of nonsense transcripts 1-like protein | 17329634 | 17332325 | chr9_15468213 | 3.07E-05 |
